# Neural Networks for parameter estimation in microstructural MRI: a study with a high-dimensional diffusion-relaxation model of white matter microstructure

**DOI:** 10.1101/2021.03.12.435163

**Authors:** João P. de Almeida Martins, Markus Nilsson, Björn Lampinen, Marco Palombo, Peter T. While, Carl-Fredrik Westin, Filip Szczepankiewicz

## Abstract

Specific features of white-matter microstructure can be investigated by using biophysical models to interpret relaxation-diffusion MRI brain data. Although more intricate models have the potential to reveal more details of the tissue, they also incur time-consuming parameter estimation that may converge to inaccurate solutions due to a prevalence of local minima in a degenerate fitting landscape. Machine-learning fitting algorithms have been proposed to accelerate the parameter estimation and increase the robustness of the attained estimates. So far, learning-based fitting approaches have been restricted to lower-dimensional microstructural models where dense sets of training data are easy to generate. Moreover, the degree to which machine learning can alleviate the degeneracy problem is poorly understood. For conventional least-squares solvers, it has been shown that degeneracy can be avoided by acquisition with optimized relaxation-diffusion-correlation protocols that include tensor-valued diffusion encoding; whether machine-learning techniques can offset these acquisition requirements remains to be tested. In this work, we employ deep neural networks to vastly accelerate the fitting of a recently introduced high-dimensional relaxation-diffusion model of tissue microstructure. We also develop strategies for assessing the accuracy and sensitivity of function fitting networks and use those strategies to explore the impact of acquisition protocol design on the performance of the network. The developed learning-based fitting pipelines were tested on relaxation-diffusion data acquired with optimized and sub-sampled acquisition protocols. We found no evidence that machine-learning algorithms can by themselves replace a careful design of the acquisition protocol or correct for a degenerate fitting landscape.

## MAIN TeXT

### 1 INTRODUCTION

Microstructure imaging aims at using diffusion MRI (dMRI) to map salient features of the tissue (Alexander et al., 2019; Nilsson et al., 2013; Novikov et al., 2019). A central goal in microstructure imaging has been to estimate volume fraction of different microstructure components (Lampinen et al., 2020; Lampinen et al., 2019; Veraart et al., 2018). Estimating volume rather than signal fractions is however challenging because it requires the simultaneous estimation of both diffusion and relaxation properties of the different compartments. This kind of inverse problem is sensitive to degeneracy issues (Jelescu et al., 2016; Lampinen et al., 2019), in which a multitude of different model parameters can describe the acquired data equally well. Parameter estimation can also be computationally slow, preventing real-time mapping. A potential solution is to employ machine learning to accelerate the parameter estimation process. However, the current literature lacks systematic descriptions of the gains and potential drawbacks of this approach, which is surprising considering the exponential increase in interest for such methods. In this work, we use neural networks to speed up the estimation process and investigate the veracity of the estimates as well as the potential for neural networks to alleviate problems that stem from degeneracy.

Neural networks and other machine learning approaches have been applied before to accelerate the estimation of microstructure parameters from dMRI data (Barbieri et al., 2020; Bertleff et al., 2017; Golkov et al., 2016; Grussu et al., 2020a; Gyori et al., 2019; Hill et al., 2021; Nedjati-Gilani et al., 2017; Palombo et al., 2020; Reisert et al., 2017). Examples include the use of a random forest regressor to compartment models with permeability for white matter microstructure imaging in presence of water exchange (Nedjati-Gilani et al., 2017) and the SANDI model to map gray matter properties (Palombo et al., 2020). Reisert *et al*. applied machine learning to a Bayesian estimation approach which dramatically accelerated the fitting of two- and three-compartment models (Reisert et al., 2017). Barbieri *et al* applied deep neural networks to the intra-voxel incoherent motion model (Barbieri et al., 2020). Nevertheless, an open question is what impact the training strategy has on the fitting performance, in particular when applied to models with many model parameters. Here, we will loosely refer to these as ‘high-dimensional models.’ For such models, generation of training data is challenging due to the poor scaling behaviour when a finite number of points are distributed across *p* parameter dimensions; to sample each combination of model parameters in *m* steps requires *m*^*p*^samples. As *p* increases, it is unavoidable that a finite set of samples becomes sparse in the *p*-dimensional space. Here, we investigate the impact that the model parameter space sampling pattern has on the performance of the neural network.

Apart from accelerating model fitting, neural networks may in principle also reduce the requirements on the imaging protocol by exploiting parameter correlations. For example, priors learned from the training data have been observed to stabilise model fitting performance against substantial degrees of data down-sampling (Alexander et al., 2017; Golkov et al., 2016; Tian et al., 2020). However, we do not expect machine learning approaches to completely alleviate degeneracy issues. Indeed, for cases where the acquisition protocol does not provide sufficient information to resolve between different parameter values, the learning-based estimates will simply equal the mean of the model parameter distribution used for training (Reisert et al., 2017).

The aim of this study was to compare training strategies, propose tools to evaluate the performance of the neural network, and test to what degree neural networks could help solve the degeneracy problem. As a testbed, we use a high-dimensional relaxation-diffusion microstructure (Lampinen et al., 2020; Lampinen et al., 2019; Veraart et al., 2018). The parameter estimation was enabled by the use of state-of-the-art imaging protocols featuring so-called b-tensor encoding (Topgaard, 2017; Westin et al., 2016), combined with diffusion-relaxation correlations (de Almeida Martins et al., 2020; de Almeida Martins and Topgaard, 2018; Lampinen et al., 2019). We also investigated if neural network-based estimation of model parameters could offset the need for tensor-valued diffusion encoding, to enable this approach for data acquired with conventional diffusion encoding.

## 2 THEORY

White matter (WM) microstructure can be modelled by multiple compartments with different micro-structural properties but a common orientation distribution (Alexander et al., 2019; Novikov et al., 2019). In this description, the measured signal is the convolution between an orientation distribution function (ODF) 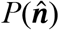 and a microstructural kernel 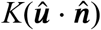

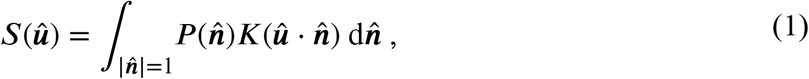

where 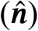 and ***û*** are unit vectors defining the symmetry axes of the ODF and of the diffusion encoding process, respectively. In this work, we assign an effective transverse relaxation time *T*_2_ and an apparent microscopic diffusion tensor **D** to each microstructural component, and use exponentially decaying functions to model the effect of these microstructural properties on the relaxation-diffusion-weighted signal (Veraart et al., 2018). Under these assumptions, the microstructure kernel is written as a weighted sum of exponentials

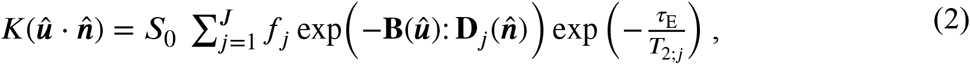

corresponding to a mixture of *J* components each with signal fraction *f*_*j*_, transverse relaxation time *T*_2;*j*_, and diffusion tensor **D**_*j*_. Information about *T*_2;*j*_ and **D**_*j*_ is encoded into the signal by the echo-time *τ*_E_ and diffusion encoding tensor **B**(***û***), respectively, both of which are experimental variables. To simplify the model, we only consider axisymmetric **B**(***û***) and additionally assume that the component-wise **D**_*j*_ are axially symmetric.

The convolution expressed in Eq. (1) can be simplified by factorizing both 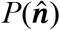 and 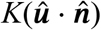 in their spherical harmonic coefficients *p*_*lm*_ and *k*_*lm*_, respectively:

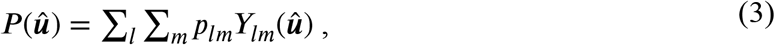

and

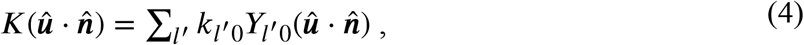

where *Y*_*lm*_ are the spherical harmonics basis functions

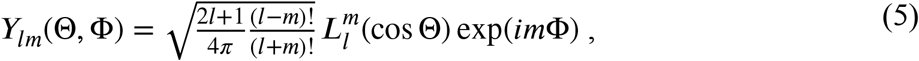

with the 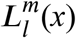 term denoting the associated Legendre polynomials. The summations in Eqs. (3) are carried out for order *l* = 0, 1, 2, …, and degree *m* = −*l*, − *l* + 1, …, *l*. In Eq. (4), we have taken the axial symmetry of the microstructural kernel 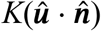 into account (Lampinen et al., 2020; Novikov et al., 2018). Symmetry around the polar axis implies *k*_*l’m’*_ = 0 for either *m’* ≠ 0 or odd *l’*. Taken together, this means that the *k*_*l’m’*_ coefficients are reduced to their 0^th^ degree terms *k*_*l’*0_ (typically written as *k*_*l’*_) and only even-ordered spherical harmonic terms (*l’* = 0, 2, …) provide non-trivial contributions. Using the spherical harmonics addition theorem, Eq. (4) can be rewritten as:

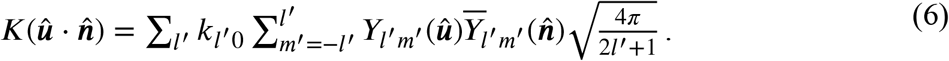

Inserting Eqs. (3) and (6) into Eq. (1) and making use of the orthogonality of the spherical harmonic basis finally yields (Driscoll and Healy, 1994; Healy et al., 1998):

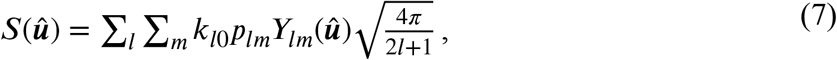

where ***û*** can be parameterized by the polar and azimuthal angles, *θ* and *ϕ*, describing the orientation of **B, *û*** ≡ (sin*θ* cos*ϕ*, sin *θ* sin *ϕ*, cos *θ*).

Exploiting the orthogonality of the spherical harmonic basis, the spherical harmonic coefficients of the ODF (*p*_*lm*_) and the microstructure kernel (*k*_*l*0_) can be determined by multiplying either 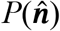 or 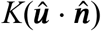, respectively, with the complex conjugate of *Y*_*lm*_ and then integrating over a sphere. For the microstructural kernel, such procedure results in:

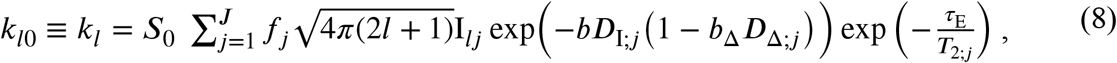

where *b* is the conventional *b*-value and *b*_Δ_ denotes the normalized anisotropy of the diffusion encoding tensor **B** (Eriksson et al., 2015). The isotropic diffusivity and the normalized diffusion anisotropy (*D*_I_ and *D*_Δ_) are related to the axial and radial diffusivities (*D*_∥_ and *D*_⊥_) of the diffusion tensor according to *D*_I_ = (*D*_∥_ + 2*D*_⊥_)/3 and *D*_Δ_ = (*D*_∥_ − *D*_⊥_)/3*D*_I_ (Conturo et al., 1996). The *I*_*lj*_ factors are a function of the regular Legendre polynomials, *L*_*l*_, and defined as:

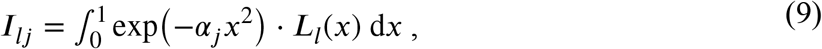

with α_*j*_ = 3*bD*_I;*j*_*b*_Δ_*D*_Δ;*j*_.

Different diffusion MRI models vary in their number of components and the constraints imposed on their properties. Here we consider a two-compartment model (*J* = 2) comprising a “stick” component (S) with zero radial diffusivity and a “zeppelin” (Z) component with *D*_Δ;Z_ ≠ 1:

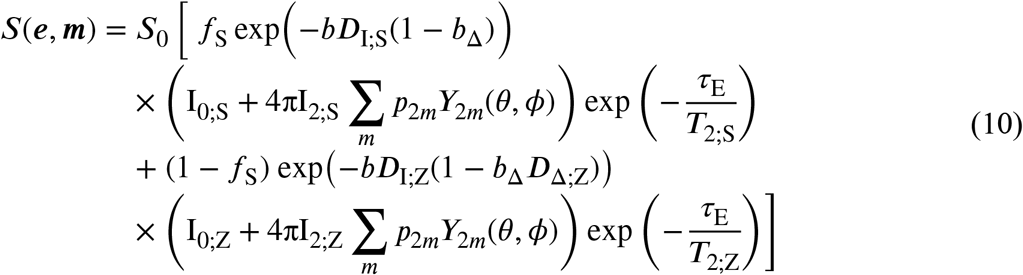

where the spherical harmonic summation is truncated at the second-ordered, *m* ∈ {−2, −1, 0, 1, 2}. The derivation of Eq. (10) uses the 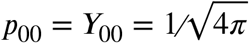 ODF normalization (Lampinen et al., 2020; Novikov et al., 2018). The vectors ***e*** and ***m*** capture the experiment-related parameters, ***e*** = (*τ*_E_, *b, b*_Δ_, *θ, ϕ*), and scalar model parameters, ***m*** = (*f*_S_, *D*_I;S_, *D*_I;Z_, *D*_Δ;Z_, *T*_2;S_, *T*_2;Z_, *p*_20_, Re(*p*_21_), Im(*p*_21_), Re(*p*_22_), Im(*p*_22_)). Besides setting *D*_Δ,S_ = 1, no other constraints were imposed on the compartment properties. We refer to the model expressed by Eq. (10) as the Relaxed Standard Model (RSM). This name is chosen to mark its descendance from the “standard model” of WM microstructure (Novikov et al., 2019) and to emphasize the fact that it accounts for compartment-specific *T*_2_ times.

The RSM model parameters can be determined by fitting Eq. (10) directly to the acquired signals (Lampinen et al., 2020). An alternative strategy, followed in (Veraart et al., 2018), is to use a model fitting framework that effectively reduces the dimensionality of the parameter space by means of performing a hierarchical factorization of the voxel-wise ODFs (Novikov et al., 2018; Reisert et al., 2017). The initial step of such framework consists in projecting the measured signal onto a spherical harmonics’ basis

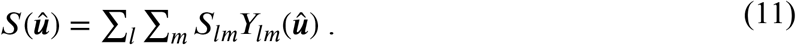

The *S*_*lm*_ coefficients are subsequently converted to rotational invariants *S*_*l*_, and fitted to the corresponding rotationally invariant terms of the *P*(**û**)⊗*K*(ĝ · **û**) convolution:

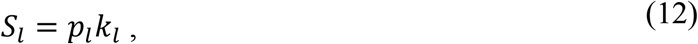

where *k*_*l*_ is the 0^th^ degree term of the microstructural kernel as defined by Eq. (8). The rotationally invariant coefficients, *S*_*l*_ and *p*_*l*_, are computed from (Novikov et al., 2018)

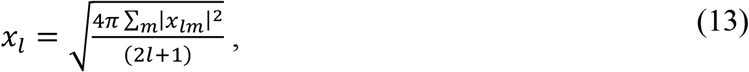

where *x*_*lm*_ are the spherical harmonic coefficients, and *x*_*l*_ ≡ *S*_*l*_ or *x*_*l*_ ≡ *p*_*l*_. Signal projections with *l* > 2 have small contributions to the measured signal (Jespersen et al., 2007), and the sum in Eq. (11) is typically truncated at the second order term (*l* = 2). The fitting framework summarized by Eqs. (11) and (12) is commonly referred to as the “RotInv” approach due to its use of rotational invariants. The *l* = 2 RotInv approach condenses the five *p*_*2m, m*∈{−2,−1,0,1,2}_ parameters of the RSM model onto a single *p*_2_ invariant capturing the orientation coherence of the sub-voxel diffusion domains, thus reducing the dimensionality of the fitting problem by four parameters.

## 3 METHODS

### 3.1 Neural network architecture and training

In this work, we constructed a feedforward deep neural network (DNN) using the *fitnet* function in MATLAB (The MathWorks, Inc.), and used it to fit vectors of scalar parameters, ***m*** = (*f*_S_, *D*_I;S_, *D*_I;Z_, *D*_Δ;Z_, *T*_2;S_, *T*_2;Z_, *p*_20_, Re(*p*_21_), Im(*p*_21_), Re(*p*_22_), Im(*p*_22_)), to sets of *S*(*τ*_E_, **B**) measurements. The network was configured to have 3 fully connected hidden layers with decreasing number of nodes (180, 80, and 55) and an output layer with 11 nodes, each of which representing a dimension within the ***m*** vectors. All hidden layers were activated by hyperbolic tangent (tanh) functions, while the output layer uses a linear activation function. The input consisted of a vector of *E* signal amplitudes sampled with a pre-defined relaxation-diffusion encoding protocol. We considered three different acquisition protocols comprising between *E* = 164 and *E* = 270 distinct (*τ*_E_, **B**) points. To remove the influence of *S*_0_ from the fitting problem, we normalized the input vector to the median of the signals measured at the point of maximal signal amplitude (minimum *b* and shortest *τ*_E_).

The choice of a fully connected DNN follows the design of classic multilayer perceptrons (MLPs), which are thus well-suited for regression problems (Cybenko, 1989; Hornik et al., 1989). In this work, we employ the tanh activation function due its stronger gradients and faster convergence (LeCun et al., 2012).

Supervised network training was performed using a scaled conjugate gradient optimiser and a mean squared error loss

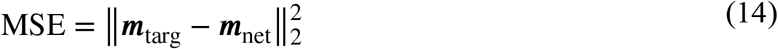

where ***m***_targ_ is the ground-truth target vector, ***m***_net_ is the corresponding network output vector, and ∥ × ∥_2_ denotes the Euclidean norm. The ***m***_targ_ parameters were rescaled between 0 and 1 using a min-max normalization strategy before being supplied to the network. The network was trained with a set of 5×10^5^ voxels with randomly generated model parameters and noisy signal *S*(*τ*_E_, **B**) (section 3.2 describes the training dataset generation). The training data was divided into different sub-sets before being supplied to the network such that 65% of the original data was used to update the weights and biases, 20% was used for cross-validation, and 15% was reserved for testing. In lieu of standard 𝓁_1_ or 𝓁_2_ regularizers, we prevented overfitting through an early stopping method and training was terminated following an increase of the MSE of the validation data for 5 consecutive epochs.

Network GPU training took approximately 3 hours on two parallel NVIDIA GeForce RTX 2080 SUPER graphic cards, each with 8 GB of memory. Both graphic cards were installed on a high-end consumer-grade desktop computer with an Intel i9-9900k 3.6 GHz CPU and 32 GB memory.

### 3.2 Generating training data

We generated training parameter vectors, ***m***_train_, from two distinct sets:

‐ parameter vectors obtained by uniform random sampling within the bounds described in **Table 1**, denoted ***m***_unif_

**Table 1.**
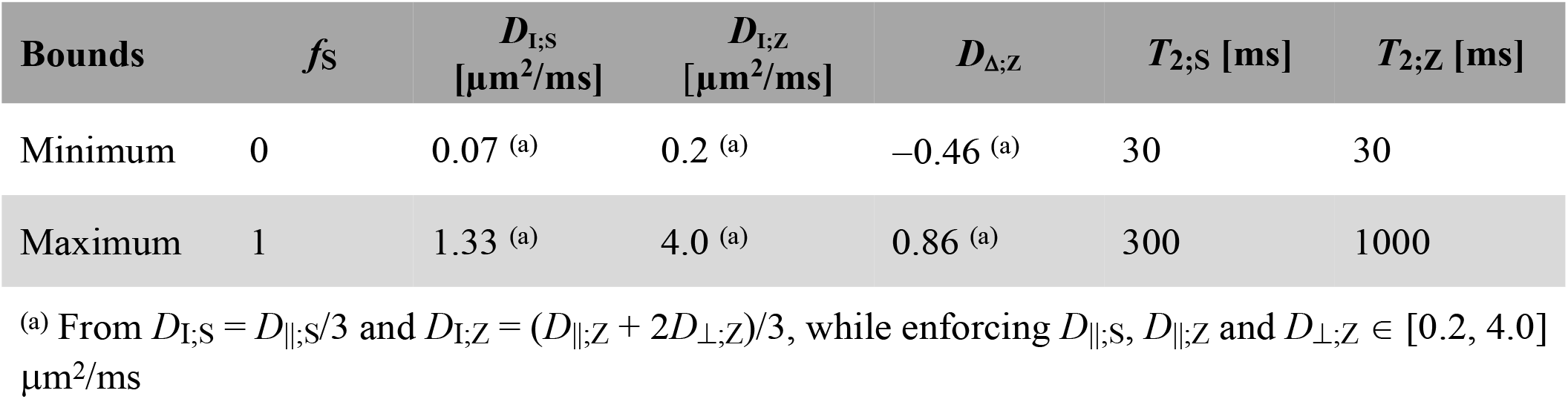
Relaxed standard model parameter bounds. The rationale behind the various model bounds is detailed in (Lampinen et al., 2020).
‐ parameter vectors estimated from a NLLS fit of Eq. (10) to *in vivo* brain data, denoted ***m***_brain_. Vectors derived from both sets were combined to create composite training datasets comprising a total of #*m*_train_ vectors:

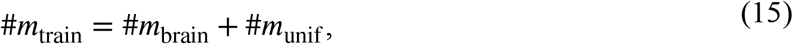

where *#m*_brain_ and *#m*_unif_ are the number of ***m***_brain_ and ***m***_unif_ vectors, respectively. To study the impact of different training data generation strategies on the accuracy of network-derived estimates, we compared the performance of networks trained with different ratios between #*m*_brain_ and #*m*_train_:

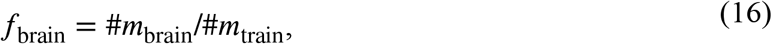

*i*.*e*., networks trained with varying relative amounts of ***m***_brain_ and ***m***_unif_ vectors. The *f*_brain_ fractions were varied between 0 and 1 in increments of 0.05, thus resulting in a total of 21 training datasets. All sets contained a total of #*m*_train_ = 5×10^5^ independent parameter vectors.

The ***m***_brain_ vectors comprise the solutions of a nonlinear least-squares (NLLS) fit of Eq. (10) to *in vivo* signal data, ***m***_fit,_ and an additional parameter set, ***m***_mut_, consisting of random mutations of the fitted solutions:

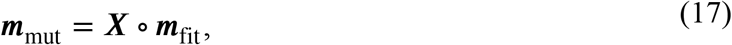

where ‘o’ denotes the element-wise (Hadamard) product, and ***X*** is an 11-dimensional vector of normally distributed numbers. Each element of ***X*** is an independent and identically distributed random variable sampled from a normal distribution with mean 1 and standard deviation 0.2. The number of ***m***_fit_ vectors was kept constant (*#m*_fit_ ∼ 8×10^4^), and the total of ***m***_mut_ vectors was defined as:

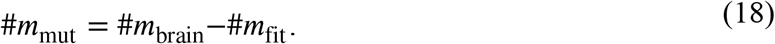

The introduction of mutated parameters is a data augmentation technique, designed to simultaneously compensate for the relative low number of ***m***_fit_ vectors and expand the (*f*_S_, *D*_I;S_, *D*_I;Z_, *D*_Δ;Z_, *T*_2;S_, *T*_2;Z_, *p*_20_, Re(*p*_21_), Im(*p*_21_), Re(*p*_22_), Im(*p*_22_)) domain of the ***m***_brain_ parameter targets.

Synthetic signal data were generated from ***m***_brain_ and ***m***_unif_ using Eq. (10) and one of three different (*τ*_E_, **B**) acquisition protocols:

‐ Protocol A comprises tensor-valued encoding with full relaxation-diffusion-correlation optimized for minimal RSM parameter variance (Lampinen et al., 2020)
‐ Protocol B comprises tensor-valued encoding with relaxation-diffusion-correlations restricted to low *b*-values(Lampinen et al., 2019)
‐ Protocol C comprises diffusion-relaxation optimized for minimal RSM parameter variance but includes only linear **B** (*b*_Δ_ = 1)(Lampinen et al., 2020).

Additional details on the various protocols can be found in their respective references and in **Table S1** of the Supporting Information.

Noise was sampled from the Rice distribution and added to the ground-truth synthetic signals. Because relaxation-diffusion MRI data displays spatially varying noise, the amplitude of the signal-to-noise ratio (SNR) was uniformly varied across voxels in the interval SNR ∈ [20, 50]. This interval mimics characteristics observed for the brain *in vivo* (Lampinen et al., 2020). Finally, networks were trained using ***m***_train_ vectors as targets and their corresponding *in silico* noisy signals as inputs.

### 3.3 Network evaluation

To find the optimal *f*_brain_ parameter, we trained networks with varying *f*_brain_, deployed them on unseen *in silico* data, and compared the various networks in terms of accuracy of the resulting parameter estimates. Network accuracy was assessed via normalized root-mean-squared errors (NRMSE) and linear correlations with ground-truth values in terms of the Pearson correlation coefficient (*ρ)*. Correlation plots were used to evaluate the network trained with the optimal *f*_brain_ value in further detail.

The effects of protocols A-C on network performance were evaluated in terms of NRMSE and sensitivity to parameter changes. The latter was gauged by modulating the parameters (*f*_S_, *D*_I;S_, *D*_I;Z_, *D*_Δ;Z_, *T*_2;S_, *T*_2;Z_) of a RSM solution, one at a time by 10%, and measuring the response in all parameters. The original parameter set was based on *in vivo* data from the *corona radiata* where *f*_S_ = 0.45, *D*_I;S_ = 0.58 *μ*m^2^/ms, *D*_I;Z_ = 1.36 *μ*m^2^/ms, *D*_Δ;Z_ = 0.44, *T*_2;S_ = 69 ms, *T*_2;Z_ = 60 ms (Lampinen et al., 2020). *In silico* datasets were subsequently generated for each of the 6 modulated datasets, noise at SNR = 100 was added to the synthetic signals, and parameter estimates were finally retrieved with protocol-specific networks.

To investigate if the reduced parameter space of RotInv fitting impacts the performance of DNN fitting, we trained a network using rotationally invariant *in silico* datasets and the same optimal *f*_brain_ value found for the RSM network. RotInv training vectors, ***m***_train;RI_, were generated from the ***m***_train_ vectors discussed in Section 3.2, using Eq. (13) to convert the full RSM parameter space to the (*f*_S_, *D*_I;S_, *D*_I;Z_, *D*_Δ;Z_, *T*_2;S_, *T*_2;Z_, *p*_2_) RotInv space. Subsequently, Eq. (12) was used to calculate *S*_*l, l*={0,2}_ signals from ***m***_train;RI_ and noise was added at SNR ∈ [20, 50] to the *in silico* data. As with the full RSM model, training was performed using the ***m***_train;RI_ vectors as targets and their corresponding synthetic noisy signals as DNN inputs. Trained RSM and RotInv networks were applied to previously unseen ***m***_unif_/***m***_unif;RI_ and ***m***_brain_/***m***_brain;RI_ synthetic datasets, and finally compared in terms of their respective target-estimate correlations.

### 3.4 *In vivo* data acquisition

We analysed data from three adult volunteers previously reported in (Lampinen et al., 2020). The study was approved by the regional ethical review board in Lund and written informed consent was obtained from all volunteers prior to scanning. Measurements were performed on a MAGNETOM Prisma 3T system (Siemens Healthcare, Erlangen, Germany) using a prototype spin-echo EPI sequence (Szczepankiewicz et al., 2019a) that facilitates user-defined gradient waveforms for diffusion encoding (Szczepankiewicz et al., 2021). Data was collected using a 2.5 mm^3^isotropic spatial-resolution, 40 slices, a matrix-size of 88×88, parallel imaging factor 2 (GRAPPA), partial Fourier of 3/4, a bandwidth = 1775 Hz/pixel, and “strong” fat saturation. Diffusion encoding was performed with gradient waveforms numerically optimized to maximize the encoding strength per unit time (Sjölund et al., 2015) and to suppress concomitant field effects (Szczepankiewicz et al., 2019b). Data was acquired for a total of 270 different combinations of *τ*_E_ and **B**, sampled according to protocol A in **Table S1** of the Supporting Information. Simultaneous multi-slice with a multiband factor of 2 was used to accelerate the acquisition (Setsompop et al., 2012), resulting in a repetition time of 3.4 s and a total acquisition time of 15 minutes.

### 3.5 *In vivo* data processing & parameter estimation

Prior to any DNN training or model fitting, all acquired data were corrected for eddy-currents and subject motion using ElastiX (Klein et al., 2009) with extrapolated target volumes (Nilsson et al., 2015). Moreover, susceptibility-induced geometric distortions were corrected using the TOPUP tool in FMRIB software library (FSL) (Smith et al., 2004), and Gibbs ringing artefact correction was performed according to the method described in (Kellner et al., 2016). To further improve the smoothness of the sought RSM parameter maps, we followed the procedure used in (Lampinen et al., 2020) and filtered the corrected data with a Gaussian kernel with a standard deviation of 0.45 times the voxel size.

The RSM model parameters were estimated from a voxel-by-voxel NLLS fit of Eq. (10) to the post-processed data. The fitting process was performed with the multidimensional dMRI toolbox (https://github.com/markus-nilsson/md-dmri) (Nilsson et al., 2018), with MATLAB’s built-in *lsqcurvefit* function being used to solve the NLLS minimization problem. To supress the frequency of outliers, model fitting was performed twice and the result with lowest residual was retained (Lampinen et al., 2020). The initial guesses were sampled uniformly from the broad parameter bounds in **Table 1**. The voxel-wise RSM solution yielding the lowest residuals was stored and used to compute *in silico* signal data following the procedure detailed in Section 3.2. In a previous study, the probability of finding the global fitting solution using two initial uniformly random guesses was estimated to approximately 99.96% for *in vivo* brain data, meaning that the final RSM solution is expected to be robust in respect to the initial random guesses (Lampinen et al., 2020). NLLS fitting of a single *in vivo* brain dataset took approximately 8 hours (∼ 5.5 s per voxel) on the CPU described in Section 3.1.

Finally, Eq.(10) was also fitted to the *in vivo* data using a DNN, which took approximately 3 seconds. Training was performed on *in silico* ***m***_train_ data with an optimal *f*_brain_ fraction. The network was trained using synthetic data generated from a single subject and deployed on the two other previously unseen subjects. Neural network fitting provided voxel-wise parameter maps that were compared to the ones obtained from a traditional NLLS fitting approach.

## 4 RESULTS

### 4.1 Impact of training set composition on network performance

We first investigated the influence of the *f*_brain_ parameter, and the result is shown in **Figure 1**. Large errors—quantified by the normalized root mean squared error (RMSE)—were found between true and estimated parameters in the WM range when the training was performed exclusively with uniformly distributed random samples (*f*_brain_ = 0). This finding can likely be attributed to the vastness of the 11-dimensional space of model parameters and that the model parameters associated with WM occupy a very small part of this space. Indeed, when drawing uniformly distributed random samples from the intervals in **Table 1** only ∼ 2% of the samples will resemble the expected properties of healthy WM tissue, i.e., *D*_I_ = *f*_S_ *D*_I;S_ + (1–*f*_S_) *D*_I;Z_ ∈ [0.8, 1.1] *μ*m^2^/ms and *T*_2_ = *f*_S_ *T*_2;S_ + (1–*f*_S_) *T*_2;Z_ ∈ [60, 120] ms. Mixing in small amounts of ***m***_brain_ led to a drastic reduction of the error, with near-constant performance obtained already for *f*_brain_ ≥ 0.1.

**Figure 1.**
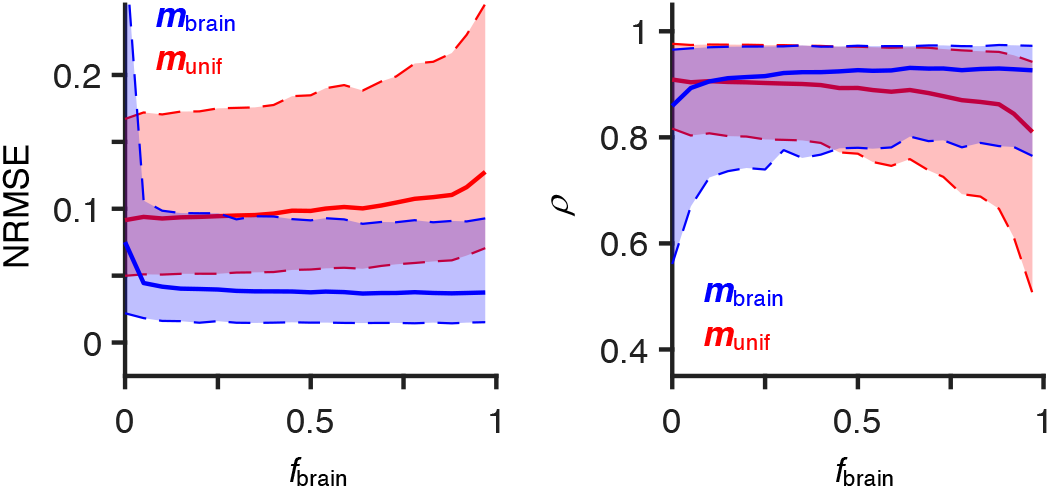
Accuracy of parameters estimated from networks trained on datasets containing varying fractions of targets derived from *in vivo* brain measurements (*f*_brain_). There is a marked trade-off between accuracy and generalizability dependent on *f*_brain_. For example, networks trained with high *f*_brain_ provide better estimates of WM-like parameters, but are constrained to a limited domain of the model space. The accuracy of network-based estimates is assessed by deploying the various networks on two previously unseen datasets where one is derived from least-squared model fitting to brain data (***m***_brain_, blue), and another obtained by uniform random sampling (***m***_unif_, red). The differences and correlations between estimated and ground-truth parameters is quantified by the normalised root mean square error (NRMSE) and Pearson’s correlation coefficient (*ρ)*. Solid lines represent the parameter-wide mean NRMSE (left plot) and the parameter-wide mean *ρ* (right plot). Computation of parameter-wide mean metrics was performed with two steps: 1) estimate NRMSE and *ρ* for each individual model parameter, and 2) average the parameter-specific metrics (NRMSE or *ρ*). The dashed lines identify the maximum and minimum estimates of parameter-specific metrics, and the shaded regions illustrate the range of metrics estimated for different model parameters.

We now turn our attention to the correlations between estimated and ground-truth parameters. For networks deployed on data derived from ***m***_brain_ vectors, higher Pearson correlation coefficients (*ρ*) are observed for higher *f*_brain_ fractions. Conversely, networks deployed on ***m***_unif_ data resulted in a monotonically decreasing relationship between *ρ* and *f*_brain_. The opposing trends are not surprising, and simply indicate that better results are obtained whenever the training and test datasets are generated with comparable strategies.

Networks trained with high *f*_brain_ fractions provide better estimates whenever the underlying data falls within the expected range of WM parameters, but high performance is constrained to a relatively small domain of model parameters. The *f*_brain_ hyper-parameter then controls a trade-off between accuracy to WM-relevant parameters and network generalizability. To achieve a balance between accuracy and generalizability, we selected the *f*_brain_ = 0.35, because it maximizes the sum between the parameter-wide mean *ρ* of ***m***_brain_ data (solid blue line in right plot of **Figure 1**) and the parameter-wide mean *ρ* of ***m***_unif_ data (solid red line in right plot of **Figure 1**). From this point onward, we concentrate on networks trained with *f*_brain_ = 0.35 datasets unless stated otherwise.

### 4.2 Neural network parameter estimates

Network-based parameter estimation was ∼10^4^ times faster than NLLS fitting on the same computer, and yielded parameters that are in good agreement with the ground-truth targets and preserve contrast between regions characterized by distinct (*T*_2_, **D**) properties (see **Figure 2**). For example, the estimated *f*_S_ and *p*_2_ maps are similar to their targets, being high in WM regions and highest in orientationally coherent WM regions such as the *corpus callosum*. An example of a slight degradation of contrast can however be observed in the *T*_2;Z_ parameter maps, where the distinction between WM (darker) and cortical GM (brighter) regions is more clear in the original map than in its NN estimate. The *T*_2;Z_ estimates are also characterized by considerable differences between ground-truth and estimated parameters in the long *T*_2_ regions such as ventricles. The largest discrepancy between estimated and target parameters was found for *D*_Δ;Z_, likely because the signal is insensitive to this parameter below values of 0.5 (Eriksson et al., 2015). Using a DNN trained on synthetic data to fit *in vivo* experimental data resulted in noisier maps that nevertheless preserve anatomically plausible contrast. The noisier appearance of the *in vivo* parameter maps is attributed to the relatively high residuals of the RSM model (Lampinen et al., 2020).

**Figure 2.**
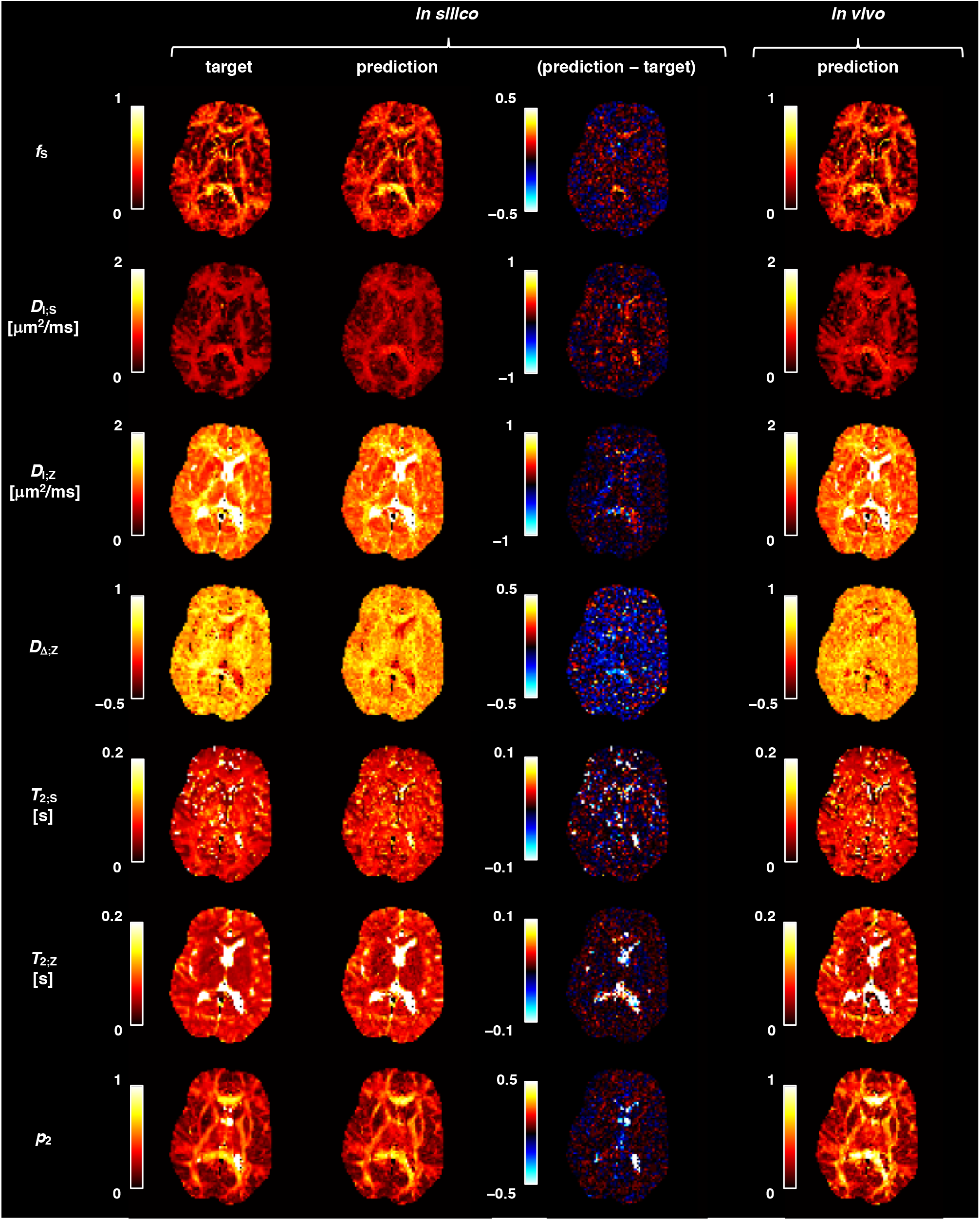
Deploying trained networks on previously unseen *in silico* and *in vivo* data provides anatomically plausible parameter maps in under 10 s (including data management times). The first and second columns compare the ground-truth targets and network predictions, respectively, of the *in silico* dataset. Difference maps are shown in the third column. Parameter maps obtained from applying a trained network on *in vivo* brain data are displayed in the fourth column.

Figure 3. shows that network-based estimates correlate well with the ground-truth parameter targets, with most parameters yielding linear correlation coefficients above 0.9. The referenced figure focuses on the performance of a network trained with *in silico S*(*τ*_E_, **B**) data generated with protocol A and *f*_vivo_ = 0.35, and distinguishes between performance on parameters obtained by uniform random sampling (light blue points) and parameters derived from *in vivo* non-cortical brain data (dark blue points). Red points correspond to parameter vectors derived from low compartment-specific signal fractions, as described in the figure caption. A considerably poor performance is observed for low *D*_Δ;Z_ values, where the network yields *D*_Δ;Z_ ∼ 0.3 regardless of the underlying ground-truth. This can be attributed to an intrinsic difficulty in distinguishing between the diffusion-weighted signals of |*D*_Δ;Z_| < 0.5 components (Eriksson et al., 2015). Moreover, weak correlations are observed for *T*_2;Z_- times much longer than the maximal *τ*_E_ of protocol A.

**Figure 3.**
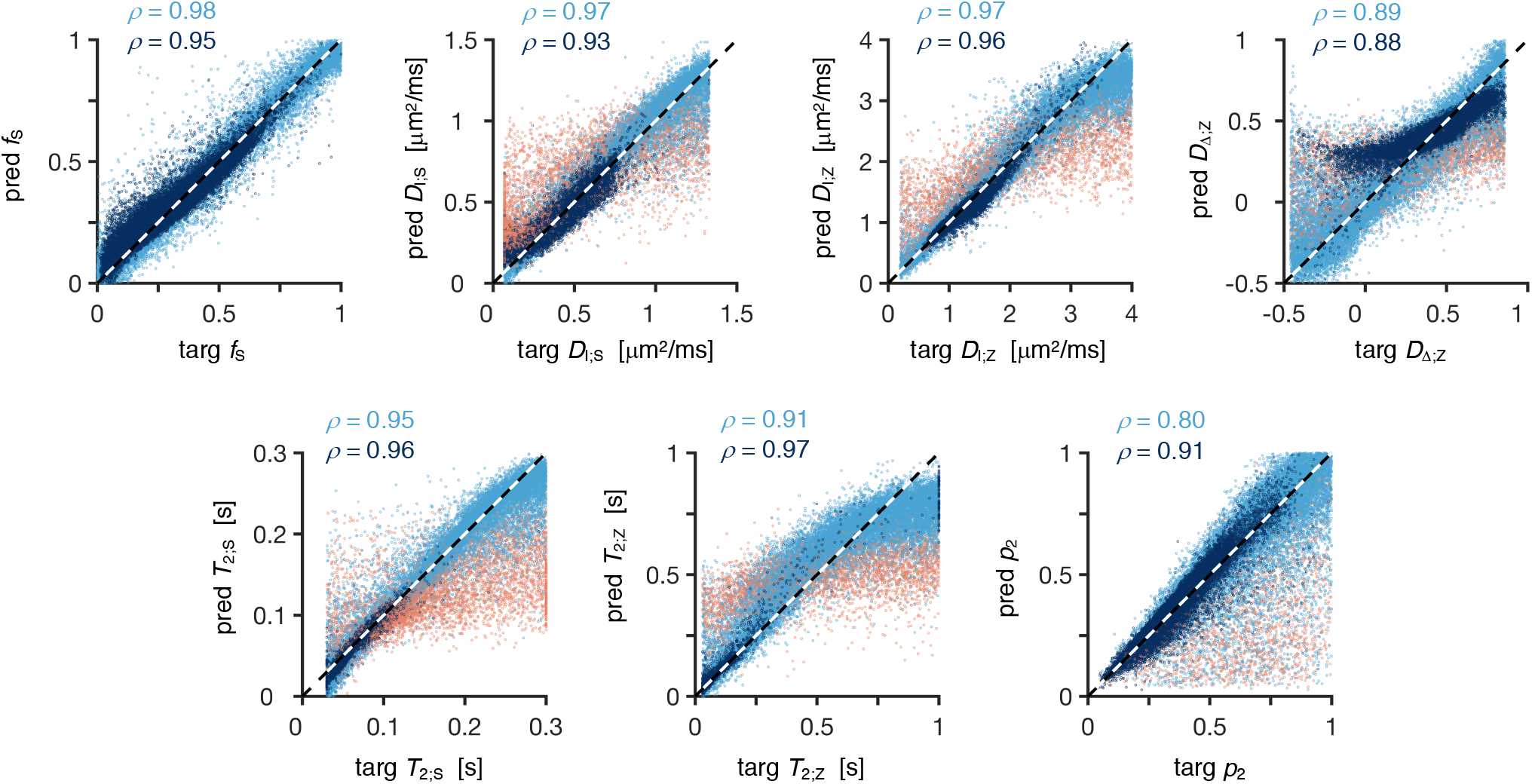
Scatter plots of ground-truth parameters vs. neural network predictions. Light blue points show results when the network is deployed on uniformly distributed random model parameters. The dark blue points correspond to an *in silico* dataset derived from a nonlinear least-squared fit to measured brain data where voxels within CSF and cortical GM were excluded by masking out regions where microscopic anisotropy (Lasič et al., 2014), *μ*FA, is lower than 0.6. The red points correspond to regions where poor accuracy is expected, i.e., where the signal fraction of the relevant component (“stick” or “zeppelin” depending on the parameter) accounts for less than 15% of the total signal or, for the *p*_2_ map, parameter vectors where the “zeppelin” component accounts for more than 85% of the total signal fraction and |*D*_Δ;Z_| < 0.4. The inner legends show the Pearson correlation coefficients (*ρ*) of the blue points.

The errors and prediction-target correlations of the network-based estimates are compiled in **Table S2** of the Supporting Information, where they are additionally compared to the errors and correlations obtained with a conventional NLLS solver. The NLLS approach has a higher accuracy for *in silico* datasets designed to capture non-cortical (*T*_2_, **D**) properties. By contrast, the function-fitting network is observed to be more accurate than the NLLS approach for synthetic ***m***_unif_ datasets.

### 4.3 Effect of acquisition protocol on network accuracy and sensitivity

In this section, we focus on the relationship between acquisition protocol design and network performance. **Figure 4** shows that network-based fitting could not improve the known fit degeneracy in protocols B and C. Indeed, networks based on the full relaxation-diffusion-correlation optimized protocol (protocol A) consistently provide lower estimation errors. Comparing protocols B and C, we note that protocol B results in a better performance when the test data is generated from ***m***_unif_ and observe a mixed performance for *in silico* test data based on ***m***_brain_ targets (protocol B yields more accurate estimates of *D*_I;S_, *D*_I;Z_, *T*_2;Z_, and *p*_2_, while protocol C yields more accurate estimates of *f*_S_, *D*_Δ;Z_, and *T*_2;S_).

**Figure 4.**
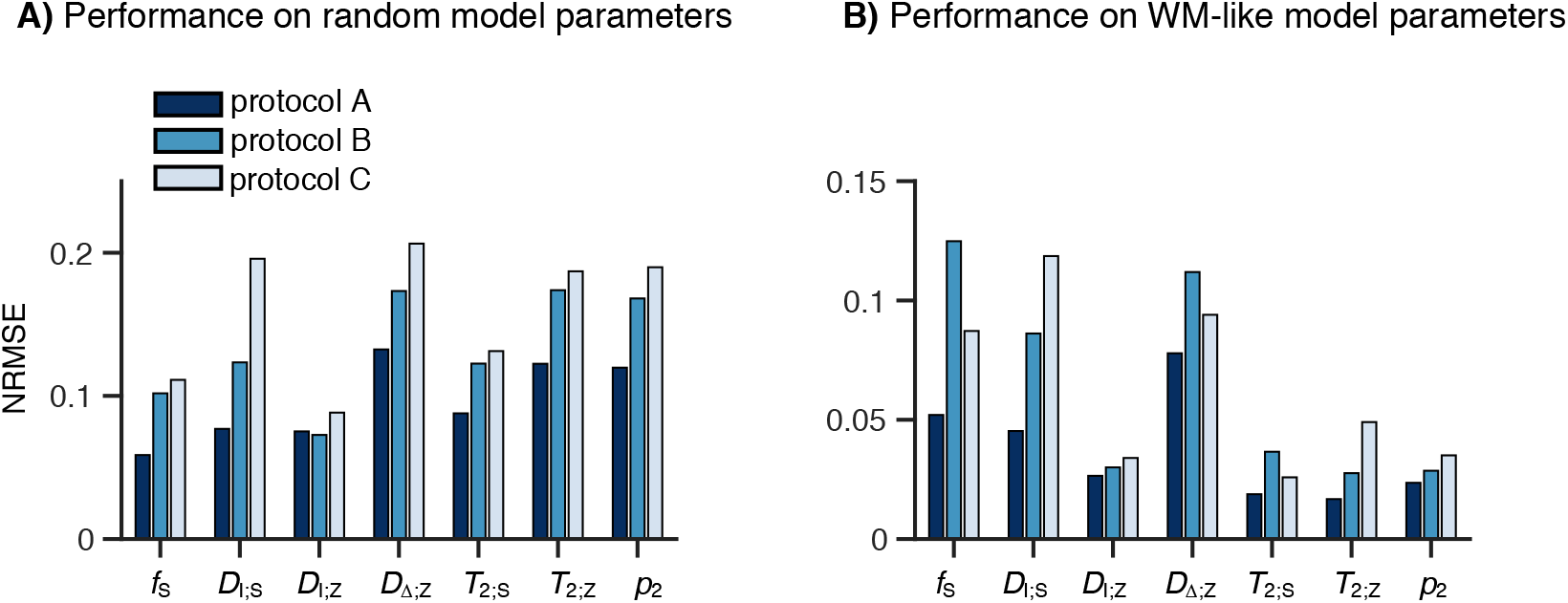
Optimized acquisition protocols result in learning-based parameter estimates with lower errors. The bar plots indicate the normalized root-mean-squared errors (NRMSE) between ground- truth and predicted parameters, for *in silico* datasets generated with different acquisition protocols. Protocol A corresponds to a tensor-valued (*τ*_E_, **B**) protocol optimized for minimal parameter variance (Lampinen et al., 2020), Protocol B is a sub-optimal tensor-valued (*τ*_E_, **B**) protocol where relaxation- diffusion correlations are exclusively established at low *b*-values (Lampinen et al., 2019), and Proto- col C is a (*τ*_E_, **B**) protocol limited to linear diffusion encoding (*b*_Δ_ = 1) and optimized for parameter precision (Lampinen et al., 2020). Panel **A** shows network performance on parameters sampled from a uniform distribution, and panel **B** shows the performance on *in silico* data based on least-squares fitting results to *in vivo* non-cortical brain tissue data.

Figure 5. shows the sensitivity of the various protocols to small parameter changes. Networks trained on data generated with protocol A are sensitive to 10% parameter modulations, but underestimate the magnitude of the change slightly. The parameter-specific modulations did not have a major effect on the estimation of the remaining unmodulated parameters. An exception occurs when the underlying T_2;Z_ is increased by 10%, which results in a 4% underestimation of the unperturbed *f*_S_. Compared to protocol A, protocols B and C exhibit a lower sensitivity to the small parameter modulations and appear to be unresponsive to changes in *D*_Δ;Z_ and *D*_I;S_, respectively. Besides its lower sensitivity, protocol C also results in less accurate estimations of the unmodulated parameters, with a 10% modulation of T_2;Z_ leading to 6% increase of the estimated *f*_S_.

**Figure 5.**
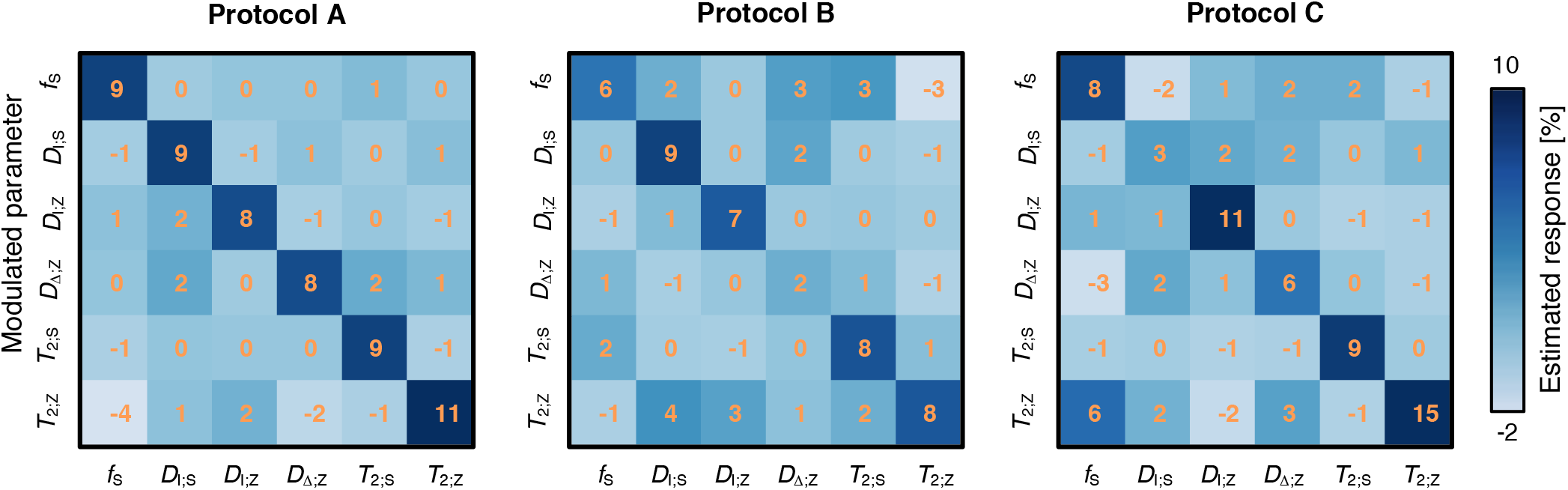
Sensitivity of acquisition protocols to 10% parameter modulations. The matrices display the relation between an induced parameter change and the observed response. When a single parameter on the y-axis is modulated by 10%, the response can be read in all other parameters along the *x*- axis. An ideal network would report a diagonal matrix with the value 10% on the diagonal, and zero otherwise. Protocol A appears sensitive in all parameters, whereas Protocols B and C lack sensitivity to *D*_Δ;Z_ and *D*_I;S_, respectively.

### 4.4 Neural network fitting of Rotationally Invariant microstructural features

**Figure 6A** displays parameter maps of *f*_S_, *D*_I;S_, *D*_I;Z_ and *D*_Δ;Z_, obtained by applying a network trained with rotational invariants to an unseen *in vivo S*_*l*={0,2}_ dataset. The resulting maps have a smooth appearance and exhibit anatomically plausible contrast. For example, regions with high *f*_S_ correspond to WM regions, the lateral ventricles are characterized by low *f*_S_ and high *D*_I;Z_ values, and darker/brighter *D*_Δ;Z_ regions demarcate cortical/non-cortical parenchyma. While it is tempting to favour the seductively ‘robust’ maps of **Figure 6A** over the noisier maps of **Figure 2** (fourth column), we note that the RotInv-based network fitting approach results in worse correlations between target and estimated parameters (compare the scatter plots of **Figure 6B** with those of **Figure 3**). For example, network-based estimates of *D*_Δ;Z_ might yield a smooth map that appeals to our intuition, but a closer look reveals that the *D*_Δ;Z_ estimates in WM and deep GM regions are equal to the mean of the target *D*_Δ;Z_ distribution and constitute a very poor estimate of the underlying ground-truth. A similarly poor correlation performance has been reported in a previous machine learning study of RotInv model fitting (Reisert et al., 2017).

**Figure 6.**
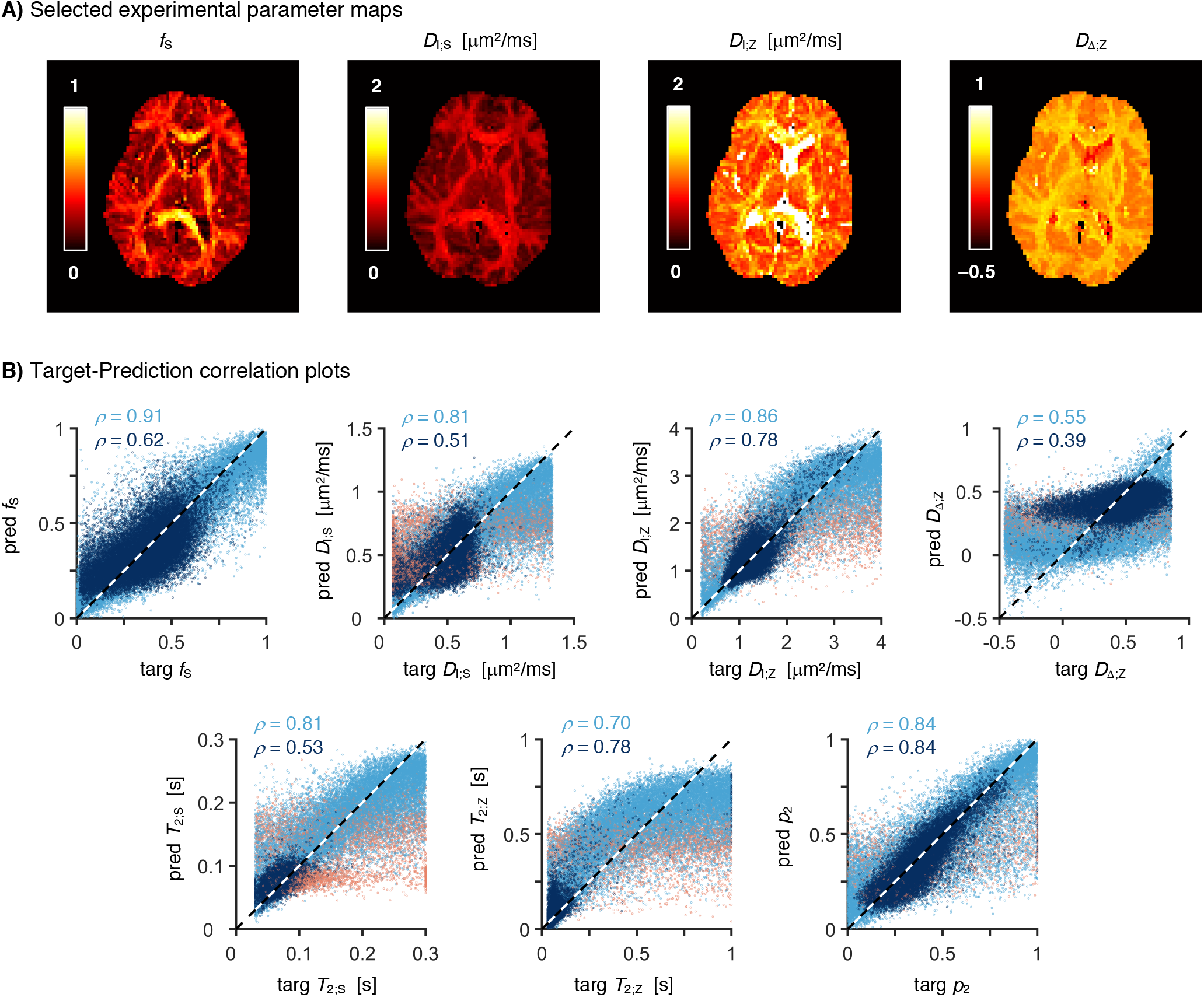
Neural network fitting of a rotationally invariant (RotInv) model results in plausible maps but poor target-estimate correlations. **A)** Maps of microstructural diffusion parameters – *f*_S_, *D*_I;S_, *D*_I;Z_ and *D*_Δ;Z_ – obtained from fitting a RotInv network to rotationally invariant *in vivo* brain data. The RotInv network was trained using a fraction of *f*_vivo_ = 0.35 between rotationally invariant ***m***_brain_ and ***m***_unif_ training parameter vectors. **B)** Correlations between network-based parameter estimates and ground-truth parameter targets. The colour-coding and legends follow the same convention as **Figure 3**.

## 5 DISCUSSION & CONCLUSIONS

Replacing traditional NLLS solvers with function-fitting neural networks provides a means to vastly reduce the time required for parameter estimation with high-dimensional microstructural models. Parameter estimation with networks trained on *in silico* data was achieved within seconds on a consumer-grade desktop computer. The resulting parameter estimates were generally observed to be in good agreement with both synthetic test datasets designed to simulate the relaxation-diffusion properties of healthy WM microstructure and test datasets spanning the entire space of allowed model parameters. When deployed on unseen *in vivo* brain data, neural networks provide maps that are consistent with known brain anatomy and preserve contrast between regions with different relaxation-diffusion properties. Our findings are encouraging and in line with recent advanced dMRI modelling studies that use machine learning techniques for parameter estimation (Barbieri et al., 2020; Bertleff et al., 2017; Golkov et al., 2016; Grussu et al., 2020a; Gyori et al., 2019; Hill et al., 2021; Nedjati-Gilani et al., 2017; Palombo et al., 2020; Reisert et al., 2017). A combination of simple error metrics, correlation analysis, and sensitivity matrices was found to provide a useful set of tools for quantitatively assessing parameter-specific accuracy/sensitivity and for identifying the limitations of learning-based approaches. These tools allowed the identification of a heterogeneous performance across the various dimensions of the RSM model, with *D*_Δ;Z_ estimates being consistently more inaccurate than other parameter estimates. However, this was unsurprising given the known difficulties of accurately estimating the anisotropy of microscopic **D** in general (Eriksson et al., 2015), and the anisotropy of the “zeppelin” compartment in particular (Lampinen et al., 2020; Lampinen et al., 2019). By contrast, simple visual inspection of machine-learned parameter maps was found to provide limited insight on the general performance of the networks. Indeed, smooth and anatomically plausible maps can be achieved even if the correlation between estimation and target is weak; a deceptive pitfall that has also been reported by (Reisert et al., 2017).

We found no evidence that learning-based fitting pipelines can by themselves navigate a degenerate fitting landscape (Jelescu et al., 2016) or replace an exhaustive and careful probing of all relevant experimental dimensions (Coelho et al., 2019; Lampinen et al., 2020). Comparison between networks based on optimal protocols and networks based on sub-optimal protocols revealed clear differences in both accuracy and sensitivity of the resulting parameter estimates. The network trained on an optimized protocol (Lampinen et al., 2020) consistently outperformed networks trained on less adequate sampling schemes. Our results suggest that the learning approach cannot substitute for a rich set of data, and learning-based fitting of diffusion-based microstructural models should be complemented with b-tensor encoding strategies (Szczepankiewicz et al., 2021) and optimized (*τ*_E_, **B**) sampling schemes. In this work, we tested the learning-based approach on protocols designed via Cramer-Rao lower-bound optimization (Alexander, 2008; Coelho et al., 2019; Lampinen et al., 2020). While not yet tested in conjunction with learning-based fitting pipelines, alternative optimization methods based on either efficient signal decomposition schemes (Bates et al., 2020; Song and Xiao, 2020) or deep learning algorithms for feature selection (Grussu et al., 2020b; Pizzolato et al., 2020) are also expected to have a positive impact on the performance of the DNN fitting approach.

The 11-dimensional parameter space of the RSM model is hard to sample densely and thus presents a challenge when designing training datasets that are representative of the vast fitting landscape. In this work, we addressed this challenge by defining composite training datasets that combine varying relative amounts of parameter vectors derived from *in vivo* healthy brain data (***m***_brain_) and parameter vectors obtained from uniform random sampling of the entire model parameter space (***m***_unif_). Networks trained exclusively with ***m***_brain_ vectors display the best accuracy in terms of expected WM properties, but their domain of validity is restricted to the relatively small space spanned by ***m***_brain_ solutions. The constrained domain of ***m***_brain_ networks raises questions about their usefulness in practical applications focusing on WM microstructural alterations, where atypical microscopic tissue structures may lead to significant deviations from the RSM parameters found in the healthy human brain (Alexander et al., 2019). To define a good trade-off between accuracy and generalizability we optimized the fraction *f*_brain_ between the number of ***m***_brain_ and ***m***_unif_. However, we expect that more work is needed to define a truly optimal strategy network training.

While the high-dimensional model parameter space introduces difficulties in the network training process, we found that directly reducing the model dimensionality through the computation of rotational invariants resulted in a reduced performance. Indeed, learning based on a RotInv framework yielded convincing parameter maps (reproducible, smooth, anatomically plausible), but closer inspection revealed both poor accuracy and sensitivity. Fitting an *l* = 0 RotInv network to “powder-averaged” diffusion-weighted data (Jespersen et al., 2013; Kaden et al., 2016; Lasič et al., 2014) resulted in a similarly poor performance. These observations suggest that the additional orientational information present in non-rotationally invariant data contributes valuable information to the learning-based fitting procedure.

A potential limitation of the present study is the relatively simple architecture of the trained networks, especially when compared to the complex high-dimensional RSM model. Given the impact of network design on the accuracy of its predictions (Isensee et al., 2020), future work should focus on a more elaborate design and thorough investigation of the effects of learning rate, network depth, optimization algorithms, and regularization approaches on network performance. Another improvement can be gained from incorporating mini-batch processing methods in the training process. This would allow training with larger datasets providing a denser sampling of the model parameter space and hopefully improve the accuracy and generalizability of the network-based estimates. Moreover, the supervised learning strategy used in this work should be compared against unsupervised deep learning strategies, which, when applied to bi-exponential modelling of low-*b* diffusion-weighted data, have been observed to provide more accurate estimates than NLLS or traditional Bayesian estimator (Barbieri et al., 2020; Kaandorp et al., 2020). Alternatives or complements to the fully-connected DNN architecture should also be explored. Promising avenues include the use of dropout (Gal and Ghahramani, 2016; Tanno et al., 2021) or deep ensemble strategies (Lakshminarayanan et al., 2016; Qin et al., 2021) as a means to derive uncertainty metrics, the use of network structures inspired by non-learning-based iterative fitting frameworks (Ye, 2017), or use denoising networks (Fadnavis et al., 2020; Wang et al., 2019) to minimize the amount of noise present in the data that is supplied to the function-fitting DNN. However, the sensitivity analysis presented here should be applied to any new fitting strategy to test specificity to change in single model parameters. Despite the clear room for improvement, we note that the estimated target-estimate correlations were stronger than those reported by Reisert *et al*. (Reisert et al., 2017), where a supervised learning in terms of a Bayesian estimator was used to fit a three-compartment diffusion model, and are equivalent to the correlations reported in more recent works focusing on machine-learning fitting of diffusion (Gyori et al., 2019; Palombo et al., 2020) and diffusion-relaxation MRI models (Grussu et al., 2020a).

In conclusion, function fitting neural networks can be used to vastly accelerate parameter estimation with high-dimensional microstructural MRI models. However, accurate estimation is achieved only if the measurement protocol samples adequate information; we found no evidence that learning-based approaches can replace the need for a rich set of data. Therefore, deep learning methodology in MRI microstructure modelling should be matched with comprehensive data acquisition. The adequacy of a given measurement protocol, in combination with a network for parameter estimation, can be evaluated by a suite of error metrics, estimate-target correlation plots, and sensitivity matrices. The learning-based fitting framework and the test tools developed herein may be used to evaluate network performance.

## CODE AVAILABILITY

Once the manuscript is accepted for publication, MATLAB code for training and deploying the networks discussed in this work will be shared in open source via a GitHub repository.

## ACKNOWLEDGMENTS

This study was financially supported by grants from the Swedish Prostate Cancer Foundation, the Swedish Research Council (2016-03443, 2020-04549), eSSENCE, and Cancerfonden. J. P. de Almeida Martins and P. T. While were supported by a grant from the Research Council of Norway (FRIPRO Researcher Project 302624) and M. Palombo by the UKRI Future Leaders Fellowship (MR/T020296/1).

