## Supporting Information for "Neural Networks for parameter estimation in microstructural MRI: a study with a high-dimensional diffusion-relaxation model of white matter microstructure"

### S.1 Relaxation-diffusion-correlation protocols

**Table S1** Relaxation-diffusion-correlation protocols evaluated in this work: a tensor-valued encoding protocol with full relaxation-diffusion-correlation optimised for parameter precision (Protocol A) (Björn Lampinen et al., 2020); a tensor-valued encoding protocol including relaxation-diffusion-correlations only at low *b*-values (Protocol B) (B. Lampinen et al., 2019); a relaxation-diffusion-correlation scheme optimised for parameter precision, but limited to linear diffusion encoding (Protocol C) (Björn Lampinen et al., 2020). The displayed table consist of a reproduction of Table 2 from (Björn Lampinen et al., 2020).

| **Protocol** | **Shell #** | ***b* [ms/μm$^2$]** | $b_\Delta$ | $\tau_E$ **[ms]** | $n_{dir}$ |
|---|---|---|---|---|---|
| **Protocol A** | 1, 2, 3 | 0.1, 1.0, 2.0 | 1 | 63 | 6, 15, 45 |
| | 4, 5, 6, 7 | 0.1, 1.0, 2.0, 5.0 | 1 | 85 | 6, 6, 15, 45 |
| | 8, 9, 10 | 0.1, 1.0, 2.0 | 1 | 130 | 30, 6, 30 |
| | 11, 12, 13 | 0.1, 2.0, 2.5 | 0.6 | 85 | 6, 15, 45 |
| **Protocol B** | 1, 2, 3, 4, 5 | 0.1, 0.5, 1.0, 1.5, 2.0 | 1 | 106 | 6, 6, 10, 16, 30 |
| | 6, 7, 8, 9, 10 | 0.1, 0.5, 1.0, 1.5, 2.0 | 0 | 106 | 6, 6, 10, 16, 30 |
| | 11, 12 | 0, 0.5 | 1 | 50 | 1, 6 |
| | 13, 14 | 0, 0.5 | 1 | 85 | 1, 6 |
| | 15, 16 | 0, 0.5 | 1 | 120 | 1, 6 |
| | 17, 18 | 0, 0.5 | 1 | 155 | 1, 6 |
| **Protocol C** | 1, 2 | 0, 0.4 | 1 | 50 | 6, 45 |
| | 3 | 0.9 | 1 | 60 | 15 |
| | 4 | 2.7 | 1 | 70 | 45 |
| | 5 | 8.9 | 1 | 85 | 45 |
| | 6 | 10 | 1 | 90 | 30 |
| | 7, 8, 9, 10 | 0, 2.1, 2.1, 2.2 | 1 | 100 | 6, 30, 10, 10 |

### S.2 Comparison between the accuracy performance of learning-based and NLLS fitting approaches

**Table S2** compiles the errors and prediction-target correlations of both network- and NLLS-based estimates. The comparison is performed using two distinct *in silico* datasets: one based on $\boldsymbol{m}_{\text{fit}}$ vectors from WM and deep GM data, and another based on $\boldsymbol{m}_{\text{unif}}$ vectors. Each dataset comprises a total of $10 \cdot 10^3$ parameter vectors (either $\boldsymbol{m}_{\text{fit}}$ or $\boldsymbol{m}_{\text{unif}}$) and their respective *in silico* signals. Rician noise with an amplitude sampled uniformly from the SNR $\in$ [20, 50] range was added to the ground-truth synthetic signals. The NLLS approach has a better accuracy performance for synthetic *in silico* datasets designed to capture non-cortical ($T_2$,**D**) properties. To the contrary, the function-fitting network is observed to outperform the NLLS approach for sets randomly sampled from the entire parameter space of the RSM model.

**Table S2** Accuracy performance of network- and NLLS-based fitting approaches. The performance is evaluated on synthetic data simulated from two different sets: uniformly sampled random parameters ($\boldsymbol{m}_{\text{unif}}$), and parameters derived from least-squared model fitting to *in vivo* WM and deep GM data ($\boldsymbol{m}_{\text{fit;WM-like}}$). NLLS solvers provide more accurate estimates of $\boldsymbol{m}_{\text{fit;WM-like}}$ data, while the neural network exhibits a better performance when the entire parameter space is considered ($\boldsymbol{m}_{\text{unif}}$).

| Metric | Dataset | Fitting method | Fitting time [s] | $f_{\text{S}}$ | $D_{\text{I;S}}$ | $D_{\text{I;S}}$ | $D_{\Delta;\text{Z}}$ | $T_{2;\text{S}}$ | $T_{2;\text{Z}}$ |
|---|---|---|---|---|---|---|---|---|---|
| NRMSE | $m_{\text{fit;WM-like}}$ | NLLS | 660 | 0.02 | 0.02 | 0.02 | 0.03 | 0.01 | 0.01 |
| | | DNN | 0.1 | 0.06 | 0.08 | 0.03 | 0.08 | 0.06 | 0.02 |
| | $m_{\text{unif}}$ | NLLS | 1855 | 0.07 | 0.17 | 0.17 | 0.19 | 0.13 | 0.17 |
| | | DNN | 0.2 | 0.06 | 0.11 | 0.10 | 0.15 | 0.13 | 0.15 |
| $\rho$ | $m_{\text{fit;WM-like}}$ | NLLS | 660 | 1 | 1 | 0.98 | 0.98 | 0.99 | 0.99 |
| | | DNN | 0.1 | 0.95 | 0.90 | 0.96 | 0.88 | 0.83 | 0.96 |
| | $m_{\text{unif}}$ | NLLS | 1855 | 0.97 | 0.85 | 0.81 | 0.76 | 0.91 | 0.85 |
| | | DNN | 0.2 | 0.98 | 0.92 | 0.94 | 0.85 | 0.90 | 0.86 |
